# Genomic analysis on broiler-associated *Clostridium perfringens* strains and caecal microbiome profiling reveals key factors linked to poultry Necrotic Enteritis

**DOI:** 10.1101/697383

**Authors:** Raymond Kiu, Joseph Brown, Harley Bedwell, Charlotte Leclaire, Shabhonam Caim, Derek Pickard, Gordon Dougan, Ronald A Dixon, Lindsay J Hall

## Abstract

**Background:** *Clostridium perfringens* is a key pathogen in poultry-associated necrotic enteritis (NE). To date there are limited Whole Genome Sequencing based studies describing broiler-associated *C. perfringens* in healthy and diseased birds. Moreover, changes in the caecal microbiome during NE is currently not well characterised. Thus, the aim of this present study was to investigate *C. perfringens* virulence factors linked to health and diseased chickens, including identifying caecal microbiota signatures associated with NE.

**Results:** We analysed 88 broiler chicken *C. perfringens* genomes (representing 66 publicly available genomes and 22 newly sequenced genomes) using different phylogenomics approaches and identified a potential hypervirulent and globally-distributed clone spanning 20-year time-frame (1993-2013). These isolates harbored a greater number of virulence genes (including toxin and collagen adhesin genes) when compared to other isolates. Further genomic analysis indicated exclusive and overabundant presence of important NE-linked toxin genes including *netB* and *tpeL* in NE-associated broiler isolates. Secondary virulence genes including *pfoA, cpb2*, and collagen adhesin genes *cna, cnaA* and *cnaD* were also enriched in the NE-linked *C. perfringens* genomes. Moreover, an environmental isolate obtained from farm animal feeds was found to encode *netB*, suggesting potential reservoirs of NetB-positive *C. perfringens* strains (toxinotype G). We also analysed caecal samples from a sub-set of 11 diseased and healthy broilers using 16S rRNA amplicon sequencing, which indicated a significant and positive correlation in genus *Clostridium* within the wider microbiota of those broilers diagnosed with NE, alongside reductions in beneficial microbiota members.

**Conclusions:** These data indicate a positive association of virulence genes including *netB, pfoA, cpb2, tpeL* and *cna* variants linked to NE-linked isolates. Potential global dissemination of specific hypervirulent lineage, coupled with distinctive microbiome profiles, highlights the need for further investigations, which will require a large worldwide sample collection from healthy and NE-associated birds.

## Background

Broiler chickens are solely bred for meat production, and represent a key global livestock asset; with an estimated annual production of 50 billion birds worldwide [1]. As broilers reach slaughter weight at a young age (4-7 weeks) they are susceptible to several welfare and infection issues. Importantly, poultry Necrotic Enteritis (NE), an inflammatory gut infection in chickens, is responsible for a loss of US$6 billion per annum in the poultry industry, with *C. perfringens* reported to be the main causative agent [2-5].

NE-associated pathologies are mainly characterised by gaseous lesions and mucosa necrosis in gas-filled distended small intestines [6]. Proposed key *C. perfringens*-associated factors linked to NE include α-toxin, and more recently NetB and TpeL (both pore-forming toxins) [7]. Other aetiological factors that have been shown to increase risk of NE include high-protein diets and environmental stressors, which may alter gut-associated microbial communities (i.e. the microbiota), host immunity, and co-infection with the poultry parasite *Eimeria* [8-10]. In addition, sub-clinical NE (SNE), which is a mild form of NE, is represented by poor growth and small intestinal ulcerative lesions and has also been associated with *C. perfringens* colonisation [7, 11].

*C. perfringens*, a ultra-rapid-growing anaerobic Gram positive pathogen, is known to harbour an arsenal of >20 toxins and has been associated with a wide range of gut diseases in animals, including poultry NE [5]. Specifically, toxin NetB is considered to be an essential *C. perfringens* virulence factor in NE pathogenesis, as determined in animal studies [2]. Expression of this pore-forming toxin has previously been reported to be higher (92%) in NE chicken *C. perfringens* isolates, as compared to healthy chickens (29%), thus supporting its role in disease progression [12]. This toxin is known to be encoded exclusively on conjugative plasmids, indicating horizontal gene transfer may play a role in dissemination to NetB-negative strains [13]. Collagen adhesin (encoded by gene *cna* and its variants) is another candidate disease determinant, which has been associated with chicken NE isolates in a recent bacterial genomic study [14].

The caecum represents the primary site for *C. perfringens* colonisation, which also contains the highest density of the chicken gut microbiota, therefore NE-induced alterations of this GI site are likely to reflect disease changes [15]. Moreover, the chicken caecal microbiome is known to play a protective role in pathogen resistance to other enteric pathogens, including *Campylobacter jejuni*, and as such intestinal microbiota disruption may impact development of *C. perfringens*-associated NE in broiler chickens, although direct biological impact is yet to be confirmed [9, 16, 17].

At present there are only two smaller scale WGS-based studies on broiler-associated *C. perfringens* [14, 18]. Thus, to further our understanding on *C. perfringens* dissemination and virulence profiles in the context of broiler-NE, we performed phylogenetics and in-depth comparative genomics on 88 chicken-associated *C. perfringens* isolates, (strains from public genome databases, alongside 22 newly isolated and sequenced strains). Moreover, it is unclear if and how the chicken caecal microbiome changes during NE development, therefore a small-scale microbiota profiling study was carried out to understand if there are any diseased-specific disturbances induced after *C. perfringens* infection.

## Results

### Phylogenetic analysis reveals a potentially important intercontinental lineage

We investigated 88 broiler-associated *C. perfringens* genomes (including 62 from NE-linked birds, 20 from healthy birds and 6 environmental isolates from broiler farms), spanning a 23-year period from 1993 to 2016 from 8 countries across European, Australasian and North American continents (Additional file 1: Table S1). Twenty-two *C. perfringens* genomes were sequenced and assembled specific to this study and the remaining isolates were publicly available. Broiler isolates Del1 and LLY_N11 were publicly available complete genomes sequenced using long-read sequencing and were included in analyses [4, 19]. A Maximum Likelihood (ML) phylogenetic tree was assembled using 88 isolates from 20194 SNPs identified from the alignment of 1810 core genes (Fig. 1). Clustering was assigned to define population structure via hierBAPS analysis; with the 88 isolates clustering into 5 major lineages.

**Fig. 1.**
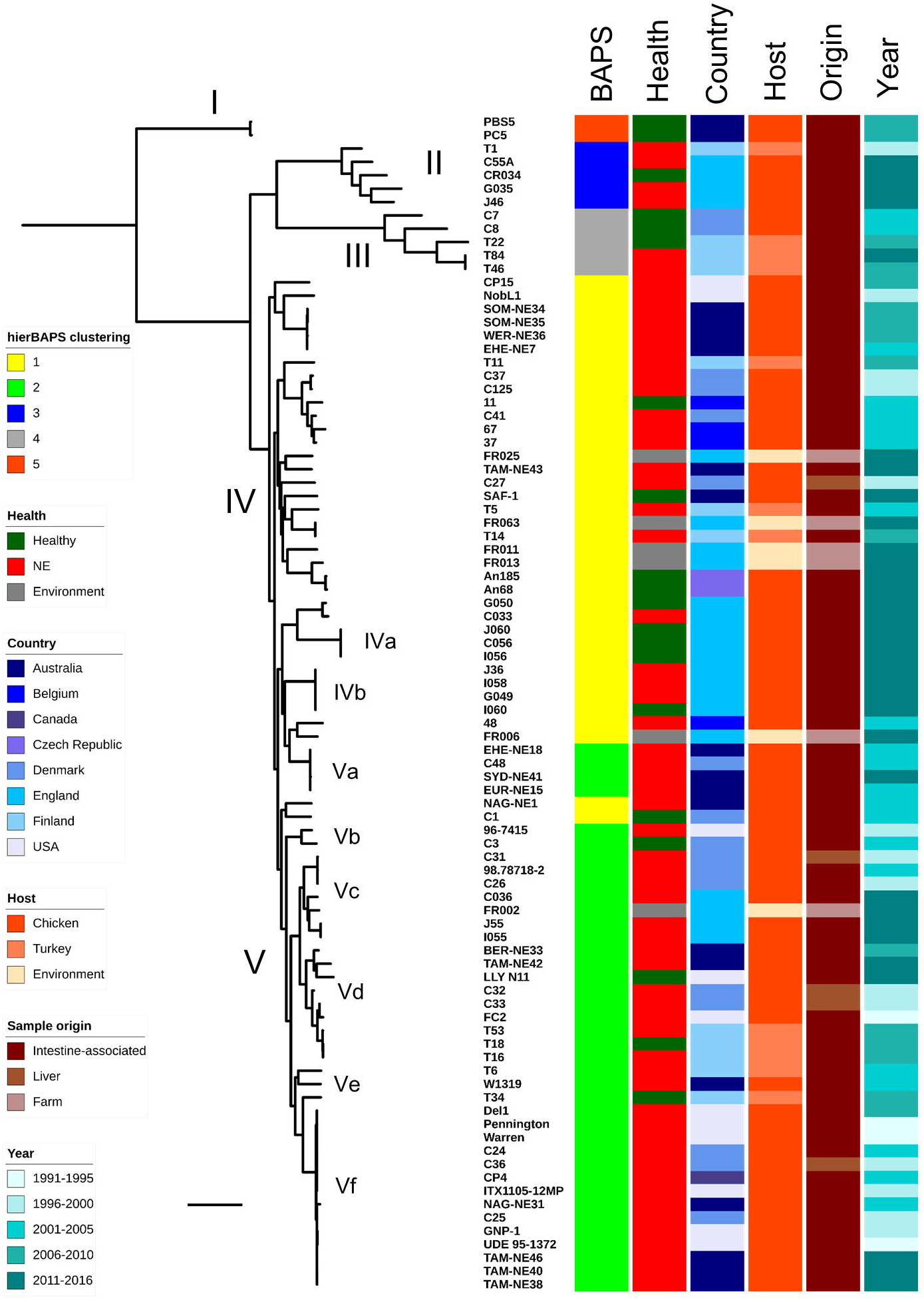
Mid-point rooted maximum-likelihood tree based on 20194 SNPs in 1810 core genes labelled by hierarchical Bayesian clusters (1-5), health states of hosts, country of isolates, poultry host species, sample origin and year of isolation. Scale bar, ∼2,000 SNPs.

The core gene alignment of 20194 SNPs was subjected to SNP distance analysis. Phylogenetic clustering and pairwise SNP analysis suggested several sub-lineages that comprised multiple highly-similar strains including lineages IVa, IVb, Va, Vc, Vd and Vf (Fig. 1). Importantly, lineage Vf that consisted of 14 broiler isolates obtained from 4 different countries namely Australia, Canada, Denmark and USA (in between 1993-2013) displayed evident clonality; 65.0 ± 52.8 SNPs between strains, which was in contrast to the closest sub-lineage Ve, with pairwise SNP distance of 1474.0 ± 162.6 SNPs (P<0.0001; Fig. 2A-C). Average Nucleotide Identity (ANI) analysis also supported the apparent clonality in sub-lineage Vf (n=14), with isolates demonstrating a pair-wise mean genome-wide ANI of 99.81 ± 0.08%, compared to closest sub-lineage pairwise ANI of 98.77 ± 1.04% (P<0.0001; Fig. 2D). Delving further into this interesting cluster of multi-continental isolates, sub-lineage Vf3 displayed the highest genetic similarity of pairwise SNP distance of 11.4 ± 6.7 SNPs;7 isolates originated from Denmark, USA and Canada, spanning a period of 16 years (Fig. 2A). Sub-lineage Vf2 displayed a similar trend of low overall pairwise SNP distance of 19.1 ± 10.6 SNPs with these 6 isolates sourced from Australia, USA and Denmark. Australian isolate NAG-NE31 was shown to be distant at 171 SNPs (Fig. 2B). This analysis suggests a potential widespread reservoir of this *C. perfringens* lineage given they are genetically highly-similar, and previous studies have indicated this species has a highly divergent genome [20-22].

**Fig. 2.**
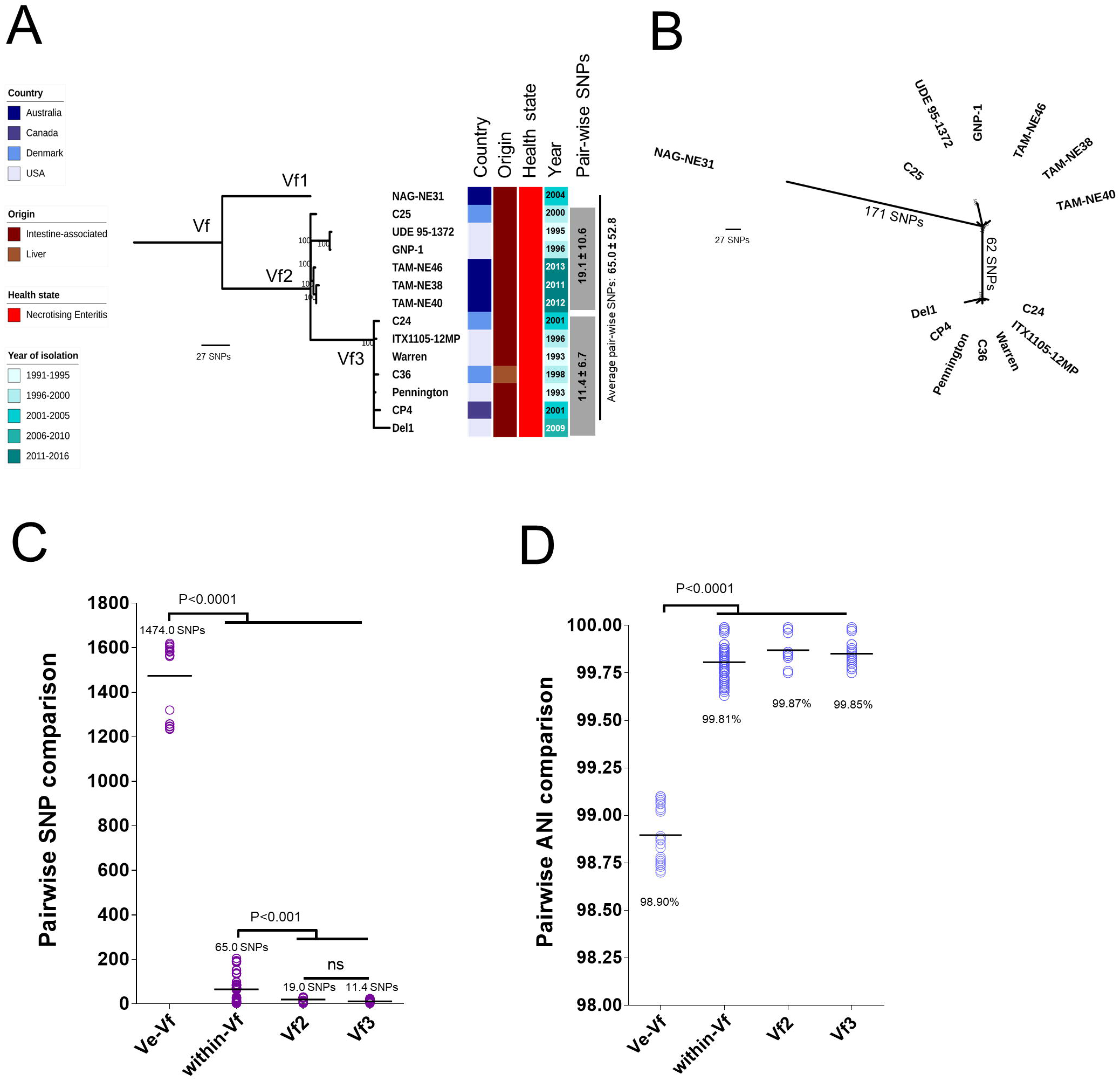
(A) Mid-point rooted maximum-likelihood tree based on 228 SNPs of 14 genomes (in 1 810 core genes) in lineage Vf, labelled by country, origin, host health state, year of isolation and within-lineage pair-wise SNP comparison. Data: Mean ± S.D. (B) Unrooted ML tree indicating SNP distance in between major branches. (C) Pair-wise SNP comparison between isolates in sub-lineages Ve vs Vf, and within sub-lineage Vf. (D) Genome wide pair-wise ANI comparison between isolates in sub-lineages Ve vs Vf, and within sub-lineage Vf. Data: Kruskal-Wallis test, Dunn multiple comparison. Means were indicated in each group.

Lineages IVa and IVb exclusively comprised newly-sequenced *C. perfringens* isolates from English farms. Lineage IVb encompassed 4 isolates from 4 individual birds (J36, I060, G049 and I058), which were identical at strain level (0 SNPs), revealing potential inter-transmission of *C. perfringens* strains among poultry farms in the same region. Interestingly, in lineage Va, Danish isolate C48 was shown to be highly similar to Australian isolates EHE-NE18 and EUR-NE15 at 7 SNPs difference, with all these isolates obtained in the same year 2002 (Additional file 2: Fig. S2). Isolates in lineages Vd and Vc exhibited geographical similarity by country at minimal SNP counts.

### Virulence association analysis supports a hypervirulent *C. perfringens* lineage

*C. perfringens* encodes an arsenal of virulence-related genes including toxin, antimicrobial resistance (AMR), and collagen adhesin genes, which are linked with gut colonisation and pathogenesis [5]. Virulence plasmids are known to encode for NE-associated toxin genes *netB* and *tpeL*. We therefore carried out a comprehensive search on all the known virulence genes, AMR determinants and virulence plasmids encoded in each *C. perfringens* genome using both assembly-based approaches (as most public genomes were only available in assemblies) and reads-mapping methods for plasmid searches (if sequencing reads available) (Fig. 3).

**Fig. 3.**
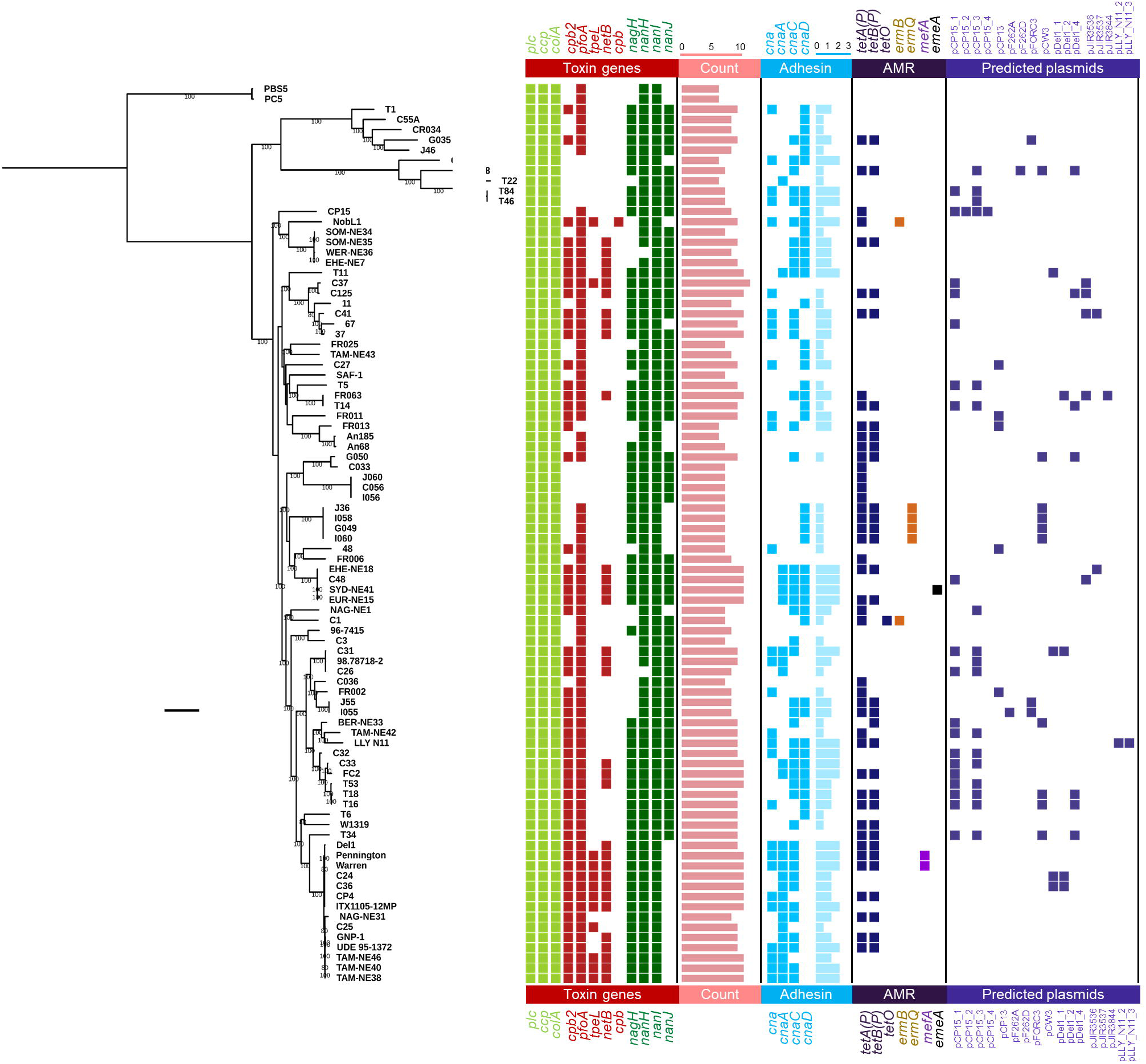
Mid-point rooted maximum-likelihood tree based on 20 194 SNPs from 1 810 core genes aligned with key virulence and resistance determinants, and predicted plasmids present in each genome for 88 broiler-associated C. *perfringens*. Coloured cells indicate predicted presence of genes/plasmids. AMR: Anti-Microbial Resistance. Scale bar, ∼2000 SNPs.

Initial analysis indicated that isolates in intercontinental lineage Vf consistently encoded more virulence genes, including *netB* and *tpeL*, thus a further comparative virulence gene analysis was performed (Additional file 2: Fig. S3). Isolates within lineage Vf encoded significantly more toxin genes (9.5 ± 0.6 toxin genes vs 8.2 ± 1.3 toxin genes; P<0.0001) and collagen adhesin genes (2.4 ± 0.6 vs 1.4 ± 1.0; P=0.0005) when compared to the remaining isolates, suggesting this is potentially a ‘hypervirulent’ sub-lineage. This was supported by further analysis comparing virulence gene counts between sub-lineage Vf, and remaining NE-linked isolates (Additional file 2: Fig. S3); Vf isolates encoded more toxin (9.5 ± 0.6 vs 8.7 ± 1.1) and collagen adhesin genes (2.4 ± 0.6 vs 1.7 ± 0.9), when compared to the remaining NE-associated *C. perfringens* strains.

Comparative analysis was also performed to define differences between NE-linked (n=62) isolates and healthy-broiler isolates (n=20; Additional file 2: Fig. S4). Both toxin genes (8.8 ± 1.1 vs 7.2 ± 1.0) and collagen adhesin genes (1.9 ± 0.9 vs 0.8 ± 1.0) were significantly elevated in NE-related isolates (P<0.0001). Considering overall virulence (toxin + adhesin genes), NE-linked isolates encoded significantly more virulence genes (10.8 ± 1.7) than healthy-broiler isolates (8.0 ± 1.6; P<0.0001). Notably, isolates in lineage Vf encode the most virulence genes (12.0 ± 1.0 genes).

To explore potential enrichment and correlation of NE-related toxin genes including *netB, tpeL* and other secondary toxin genes *pfoA* and *cpb2*, an association statistical analysis (Chi-square test) was performed (Additional file 2: Fig. S5). Toxin gene *netB* was exclusively encoded in NE-linked isolates (31/62; 50%) compared to healthy-broiler isolates (0/20; P<0.0001). Moreover, *tpeL* (12/50; 19.3%; P=0.0332), *pfoA* (59/62; 95.1%; P=0.0017) and *cpb2* (49/62; 79.0%; P<0.0001) were all shown to be enriched in NE-linked isolates. Most isolates in lineage Vf encoded these 4 toxin genes including *netB* (12/14; 85.7%), *tpeL* (10/14; 71.4%), *pfoA* (14/14; 100%) and *cpb2* (14/14; 100%), supporting the hypothesis of a hypervirulent clone.

Genome-wide association analysis highlighted additional factors that may correlate with widespread nature of lineage Vf isolates. Aside from the associations of toxin genes *tpeL* (sensitivity: 64.2%; specificity: 97.3%; 9/14 isolates) and *netB* (sensitivity: 85.7%; specificity: 72.9%; 12/14 isolates) as described above, collagen adhesin *cnaA* (sensitivity: 100%; specificity: 85.1%; 14/14 isolates) and a pilin-associated gene (group_5443; sensitivity: 100%; specificity: 86.5%; all 14/14 isolates) were specifically associated with this lineage of isolates (Additional file 1: Table S10). When we further compared the representative pilin-associated gene group_5443 using NCBI non-redundant (nr) nucleotide database via BLASTn, this gene was detected in both reference chicken isolates EHE-NE18 and Del1 complete genomes at 100% identity, which was suggested as a hypothetical protein in the annotated file (Additional file 1: Table S10). Other lineage Vf-associated genes including ABC transporter-related genes (n=4; group_1636, group_3194, group_2785, group_3195; sensitivity: 100%; specificity: 82.4%) and phage-associated genes including capsid protein (group_1646), phage-regulatory protein (group_6371) and endolysin (group_4126) were also identified.

Adhesin is an important virulence factor in broiler-linked NE [14, 18], and in this study we found that adhesin genes (at least one variant) were overall enriched (P<0.0001) in NE-linked isolates (58/62; 93.5%) vs healthy isolates (9/20; 45%). Among all related adhesin variants, *cna, cnaA* and *cnaD* genes were significantly enriched in NE-associated *C. perfringens* isolates (P<0.05), linking these genes to NE-disease development (Additional file 2: Fig. S6).

Importantly, environmental isolates encoded comparable virulence gene profiles, suggesting potential reservoir including soil and feeds. Indeed, environmental isolate *C. perfringens* FR063 was found to encode the NE-linked *netB*.

Previous studies [20, 23] have indicated that acquired AMR genes are not widespread, and a total of 7 AMR genes were detected across 88 isolates (Fig. 3). Tetracycline resistance genes *tetA(P)* and *tetB(P)* were encoded in the greatest number of genomes (44 and 32 isolates respectively), with erythromycin-resistance genes *ermB* and *ermQ* encoded in 2 and 4 isolates respectively. Macrolide-resistant efflux-pump gene *mefA* was detected in two sub-lineage Vf isolates, while multidrug-resistant gene *emeA* was detected in one isolate SYD-NE41 [24]. Notably, approximately half (47.7%) of healthy-broiler and NE-linked isolates (n=42) did not carry any acquired AMR genes.

The presence of plasmid(s) was predicted in all genomes using a reference-based sequence-search approach. Overall, 43 out of 88 (∼48.8%) isolates carried at least 1 plasmid (18 isolates carried 1 plasmid, 15 isolates harboured 2 plasmids, 4 isolates harboured 3 plasmids, 6 isolates carried 4 plasmids; detailed in Additional file 1: Table S11). Geographical association analysis indicated that two specific plasmids were present in birds from Europe, Australia and North America - plasmids pCP15_1 and pCP15_2 which were first identified in isolate CP15 from an NE-linked chicken in USA (Additional file 2: Fig. S7). These two reference plasmids did not carry any well-studied virulence-related genes, nevertheless, the re-annotation of plasmid genes using genus-specific database indicated that this small 14-kb plasmid pCP15_2 encoded a number of genes associated with sugar metabolism including phosphotransferase system (PTS), and sugar transporters sub-units (n=5; Additional file 1: Table S12). In terms of plasmid types, European isolates carried 13 different types of isolates, Australian 5 types and USA 6 types. Australian isolates were not found to encode a ‘continent-specific’, plasmid type, while Europe had 9 unique plasmid types. Plasmids pDel1_4 and pCW3 were the common plasmids detected in isolates from England, Finland and Denmark (Additional file 2: Fig. S8). Importantly, both plasmids pDel1_4 (Additional file 1: Table S13) and pCW3 (Additional file 1: Table S14) belonged to conjugative plasmid pCW3 family, carrying AMR genes *tetA(P)* and t*etB(P)* and adhesin gene *cnaC*; sharing highly similar genomic characteristics including plasmid size (47-49 kb) and CDS number (50-55; Additional file 1: Table S15)[25].

### Specific microbiota signatures identified in broiler caecal contents

To further our understanding of the broiler microbiota, particularly in the context of NE disease development, we obtained 11 caecal content samples from 11 individual broilers representing; 3 NE birds, 3 healthy birds and 5 sub-clinical NE birds.

PCA did not indicate distinct clustering of samples; suggesting a lack of distinctive microbiota signatures between diseased and healthy broilers; however healthy caecal samples appeared to have an inverse relationship with *Enterococcus* (Additional file 2: Fig. S9). Notably, disease-specific profiles might be masked by the fact that a probiotic mix was given to these broilers as a preventative measure against NE development. Therefore, raw reads from these genera were removed and another PCA was performed to understand the impact of other secondary or low abundance microbiota members. Again, health vs. disease-status clustering was not observed, however secondary NE-associated profiles did appear to positively correlate with genus *Clostridium*. Clustering at family level also did not indicate specific health-status signatures. Diversity analyses (including Inverse Simpson index, Shannon-weaver index and Fisher index) indicated there was no significant difference (P>0.05; ANOVA) in genus diversity between groups (Additional file 2: Fig. S10).

Relative bacterial genus abundance in each caecal sample was constructed to visualise microbiota profiles (Fig. 5). Thirty-seven genera were represented, with *Bifidobacterium* and *Lactobacillus* most abundant, which likely reflected the probiotic supplementation in the chicken feed. A number of secondary genera, which are usual intestinal microbiota members, were detected in these samples (relative abundance <10% in each sample) including *Blautia, Coprococcus, Dorea* and *Oscillospira. Blautia* was more abundant in health-associated caecal microbiomes (mean abundance: 3.06 ± 2.84 %) compared to diseased-associated caecal microbiomes NE (0.72 ± 0.5%) and SNE (0.14 ± 0.89%; Fig. 5B-D). The microbiota member *Enterococcus*, which is widely used as veterinary probiotic (especially *Enterococcus faecium*), was found at high levels in broilers NE2 (50.3%) and SNE5 (31.0%). Notably, *Blautia, Dorea, Oscillospira, Faecalibacterium, Coprobacillus* and *Ruminococcus* were present in all broiler microbiotas, albeit some in low abundance. Certain genera appeared to be more abundant in disease-linked NE and SNE samples including *Enterococcus* and *Bacteroides*. While on family level, Enterobacteriaceae and Enterococcaceae appeared to be elevated in NE and SNE samples. Likewise, on phylum level, phyla Proteobacteria and Bacteroidetes seem to be low-abundant in healthy samples (Additional file 2: Fig. S11).

**Fig. 4.**
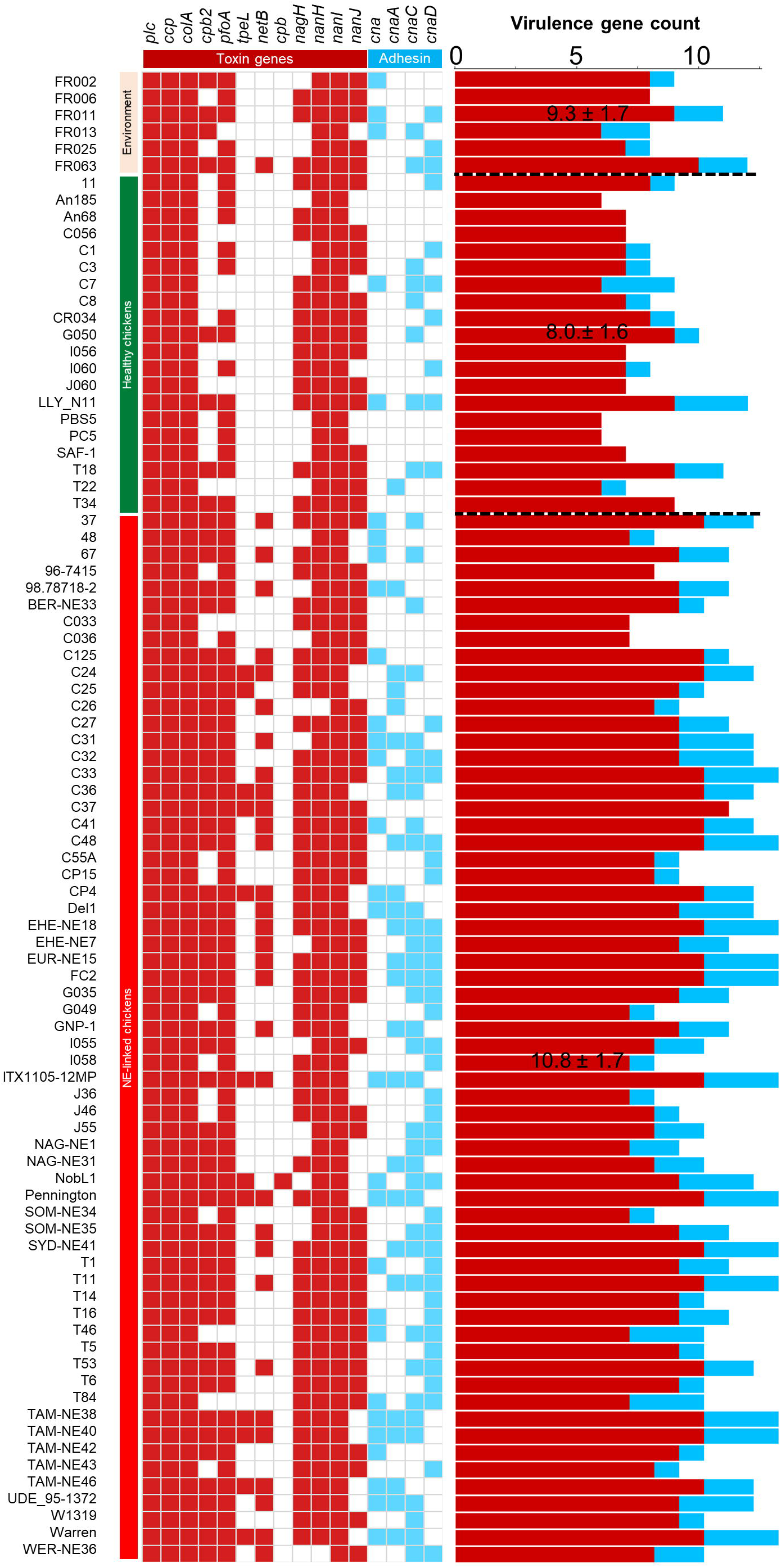
Virulence profiles of toxin genes and collagen adhesin genes categorised by host health states of bacterial isolates. Coloured cells indicate presence of gene. Mean ± S.D. in virulence gene count include toxin (red) and adhesin (lightblue) genes.

**Fig. 5.**
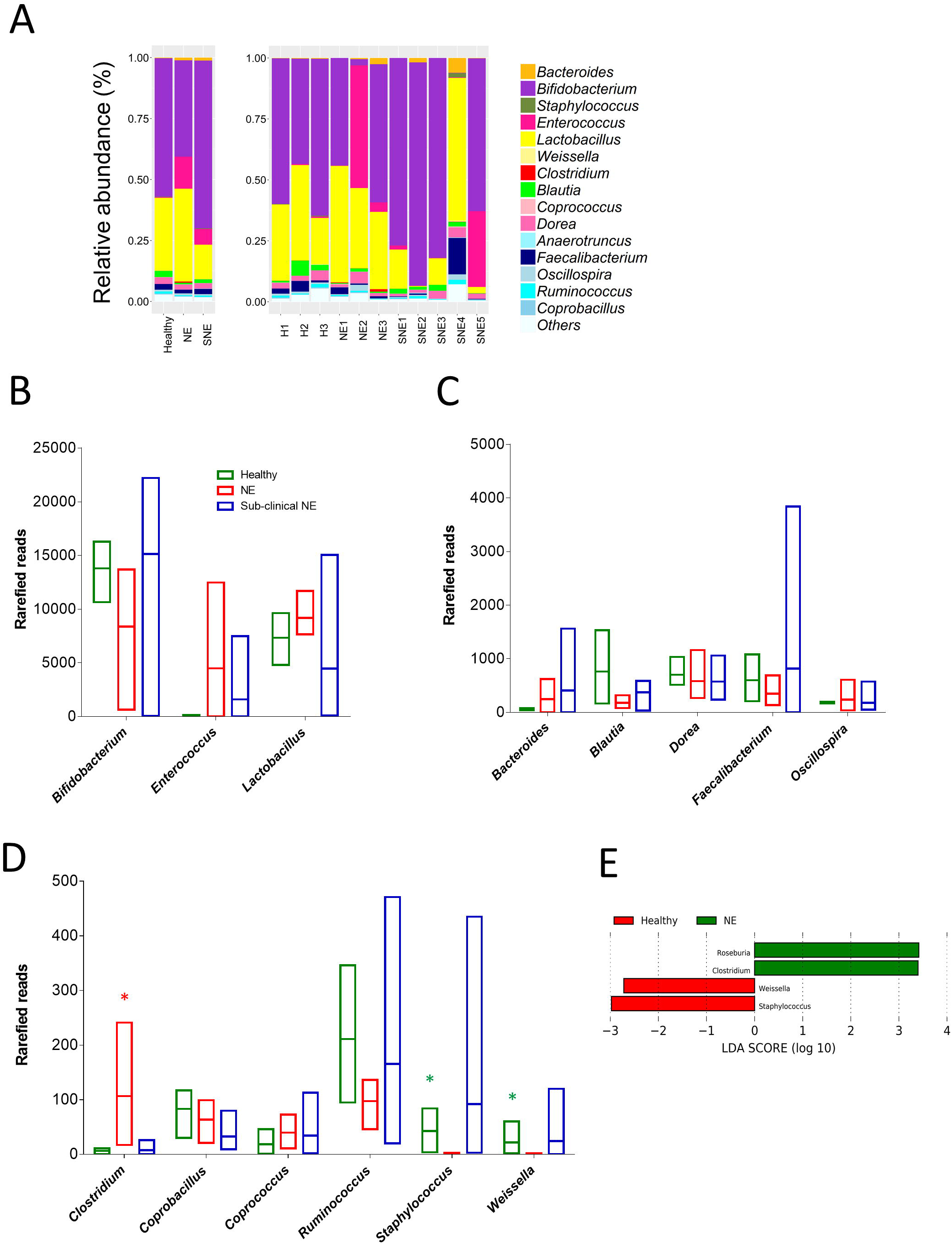
(A) Relative abundance of Operational Taxonomical Units (OTUs) based on 16S rRNA amplicon sequencing in 11 individual caecal samples on genus level, with classified comparisons on healthy, NE and SNE(sub-clinical NE) samples. (B) to (D) Comparison of taxa reads in 3 groups of samples. (E) Linear Discriminant Analysis (LOA) indicating significantly enriched taxa.

Importantly, LDA analysis indicated that *Clostridium* was significantly enriched within NE caecal microbiomes (mean relative abundance: 0.44% vs 0.03% in healthy individuals), confirming the frequent link of *Clostridium*, especially *C. perfringens*, to chicken NE. Other genera were also associated with health status: *Roseburia, Staphylococcus* and *Weisella* (Fig. 5E).

An additional paired-end BLASTn approach, to assign species level 16S rRNA sequences, indicated several important genera were present (Additional file 2: Fig. S12) [26]. *reuteri* (common broiler gut member, also widely used as probiotic supplement), *Lactobacillus salivarius* (common swine gut microbiota member used as broiler probiotic that improves production and general health) and *Lactobacillus vaginalis* (frequently found in broiler gut and a persistent gut coloniser) were the main species within the *Lactobacillus* genus [27-30]. *Enterococcus* genus primarily consisted of species *Enterococcus faecium*, a widely-used probiotic reported to promote broiler growth and suppress *C. jejuni* and *C. perfringens* infections, while stimulating the growth of *Lactobacillus* and *Bifidobacterium* [31, 32]. Importantly, *Clostridium* genus was mainly assigned to *C. perfringens* sequences, denoting the potential NE-link of *C. perfringens* origins in NE-broilers particularly NE3, where *C. perfringens* strains C036 and J36 were isolated from the same NE bird.

## Discussion

*Clostridium*, particularly *C. perfringens*, is consistently described as the primary infectious agent to cause chicken NE. As *C. perfringens* thrives at ambient bird body temperature (i.e. 40-42°C), with a doubling time <8 mins *in vitro* (the shortest generation time known for a microorganism), this may link to its ability to rapidly overgrow and cause disease pathology [5, 33]. In this study, we profiled the genomes of *C. perfringens* isolates, including newly sequenced strains, across a geographically diverse and varied health status sample collection. Genome-wide analysis revealed positive associations of important toxin genes with broiler-NE, and we identified a globally-disseminated potentially hypervirulent lineage Vf, which comprised isolates encoding important toxin genes *netB* and *tpeL* [13, 34].

*In silico* toxin profiling indicated that the NetB toxin, which has been identified as an essential toxin in NE development, [13, 34], was present in ∼50% of the NE isolates, with environmental samples also encoding this toxin, which may act as potential reservoirs, linked to NE outbreaks [35]. The fact that *netB* gene was exclusively encoded in NE-linked broiler isolates, when compared to healthy isolates, further supported the strong association of this toxin and NE pathogenesis.

Other virulence factors have also been implicated in NE pathogenesis. Several studies have indicated that collagen adhesin (encoded by *cna*) [14, 18, 36, 37] may facilitate bacterial colonisation within the chicken gut. Our analysis indicated this gene (including its variants *cnaA* and *cnaC*) was overabundant in NE-associated isolates when compared to healthy-broiler isolates (P<0.05), which also suggests a positive association with NE outcomes [37].

*C. perfringens* encodes a diverse array of toxins, and interestingly we also observed that several other accessory toxins were enriched in NE isolates, indicating these may also play an underrated role in broiler NE [38]; PFO, a pore-forming toxin which has been linked with bovine haemorrhagic enteritis [39], and CPB2, or beta2-toxin, another pore-forming cytolytic toxin associated with NE in piglets and enterocolitis in foals [40].

This genomic study indicates a potentially prevalent hypervirulent lineage Vf (comprised 14 isogenic strains; pairwise mean SNPs: 65 in 1 810 core genes), with strains obtained from Australia, Canada, Denmark and USA, spanning a period of 20 years (1993-2013). Previous analysis with 9 isolates (out of 14) also indicated these (isogenic) strains grouped within the same lineage [18]. Notably, lineage Vf isolates carried significantly more virulence genes (toxins, including *netB* and *tpeL* and collagen adhesin) than isolates in other NE-linked isolates, toxin genes, supports that this lineage may be hypervirulent. TpeL toxin is not typically considered essential for pathogenesis due to its low carriage rate among NE-linked *C. perfringens* isolates (in this study *tpeL* was exclusively detected in lineage Vf) [41]. Nevertheless, in a broiler-NE infection model, infection with *tpeL-*positive (also *netB-* positive) strains induced disease symptoms more rapidly, and with a higher fatality rate, in contrast to *tpeL*-negative strains encoding only *netB*, highlighting a role for TpeL in more severe chicken-NE pathogenesis [42]. These data also indicate a potential global dissemination of NE-associated virulent genotypes, which is in agreement with a previous study that indicated clonal expansion of *C. perfringens* via multiple-locus variable-number tandem repeat analysis (n=328)[43]. However, significantly larger sample sizes from various geographical origins will be required for in-depth WGS population structure analysis, if we are to understand the spread of *C. perfringens* in chicken farms worldwide, which will be vital in the context of disease control.

Key *C. perfringens* virulence factors including toxin and AMR genes are known to be carried on plasmids [44], including the poultry-NE-related toxin *netB* [13, 34]. The universal *tcp* conjugative system in majority of plasmids may facilitate horizontal gene transfer and enhance the virulence of *C. perfringens* strains [45, 46]. As almost half of the genomes carried plasmids (∼48.8%) this implies widespread plasmid transfer within broiler-associated *C. perfringens* strains. However, as our analysis was carried out using reference-based approaches, in some cases, fragmented short-read sequenced genome assemblies from public databases this may not readily identify plasmid sequences. Indeed, within lineage Vf we did not observe the expected high carriage rates of plasmids encoding *netB* [47]. The availability of long-read sequencing (e.g. PacBio and Nanopore) will improve investigations into *C. perfringens*, as plasmids can be sequenced and predicted more accurately despite encoding numerous tandem repeats [48, 49].

In this study, we also analysed *C. perfringens* isolates obtained from healthy or asymptomatic birds, with several isolates (LLY_N11 and T18) encoding comparable numbers of virulence genes when compared to broiler-NE isolates (n>10). Importantly, healthy-broiler isolate LLY_N11 (*netB-*negative strain, encoded *pfoA* and *cpb2*) has previously been shown to successfully induced NE in an experimental model [4, 50]. These data highlight the important role other host factors that may play a role in prevention of overt disease e.g. the chicken gut microbiota. Gut-associated microbial ecosystems are known to play a key colonisation resistance role, preventing overgrowth of so called pathobionts, or infection by known enteric pathogens (e.g. *Salmonella*).

In this small-scale broiler microbiome study, healthy broiler caecal microbiomes appeared to have enhanced abundance of the genera *Blautia*. Members of the *Blautia* genus are known to be butyrate producers, and reductions in this genus have previously been associated with a *C. jejuni* infection model [51]. Moreover, *Blautia obeum* (later reclassified as *Ruminococcus*) has also been demonstrated to restrict the colonisation of gut pathogen *Vibrio cholerae* [52, 53]. As butyrate is an important energy source for intestinal cells, these *Blautia* spp. may act as key beneficial microbiota members, serving to enhance intestinal health of chickens by strengthening the epithelial barrier, thus preventing pathogenic microbes successfully colonising and initiating disease.

In NE caecal samples we observed appearance of the *Clostridium* genus, which was significantly enriched, albeit at low reads in NE individuals. Further species-level assignment analysis indicated that most *Clostridium* sequences mapped to *C. perfringens*, indicating that even a small proportion (mean relative abundance: 0.44%) of *C. perfringens* could potentially be lethal to broiler hosts. Therefore, microbiota profiling of *Clostridium* may be useful as a potential biomarker for NE-onset, however larger studies would be required to verify these findings.

Probiotics, including *Bifidobacterium* and *Lactobacillus*, and also *Enterococcus*, have been frequently used in broiler farming primarily for growth-promotion and prevention of bacterial infections [29, 54, 55]. These taxa of beneficial bacteria have been reflected in caecal microbiome analysis, with predominant OTU proportions been assigned to *Bifidobacterium* and *Lactobacillus* across all samples. A previous study identified specific antibacterial peptides produced by *Bifidobacterium longum* that may correlate with the proposed probiotic/pathogen-inhibitory effect against *C. perfringens* [56]. Nevertheless, our analysis does not definitively verify the colonisation potential of these probiotic-associated genera in broilers’ intestines, or whether the high levels were more transient in nature, as it is common practice in poultry farms to administrate these strains in large amounts within the feed. Moreover, although there were no significant changes in the OTU proportions of these two probiotics across three groups, several birds did present with SNE and NE suggesting these strains may not be effective in reducing the disease burden associated with *C. perfringens*. However, large scale-controlled supplementation trails would need to be completed to provide robust evidence for health promotion using probiotics in poultry.

## Conclusions

In conclusion, genomic analysis of 88 broiler-associated *C. perfringens* isolates indicates positive correlations relating to virulence genes including *netB, pfo, cpb2, tpeL* and *cna* variants linked to NE-linked isolates. Furthermore, potential global dissemination of hypervirulent lineage Vf *C. perfringens* strains highlights the need for further investigations, which will require a large worldwide dataset on NE-related *C. perfringens* isolates.

## Methods

### Sample collection and bacterial isolation

Birds (culled as part of routine farm surveillance) were collected from sites reporting both healthy flocks and flocks that had been diagnosed with NE. Birds were necropsied and putative disease identification performed, followed by caecum content collection. Isolation of *C. perfringens* was carried out by isolating organs and submerging 0.1% peptone water (Oxoid, UK) in a 1:10 ratio of organ to peptone. Samples were streaked onto egg yolk agar supplemented with cycloserine (Oxoid, UK) and incubated overnight anaerobically at 37°C [57]. Single black colonies were re-streaked on brain heart infusion agar (Oxoid, UK) and incubated anaerobically at 37°C overnight. Several colonies were collected and subjected to identification of the *plc* gene by PCR, followed by 16S rRNA full-length amplicon sequencing as described previously for species verification [58, 59].

### Bacterial isolates and DNA sequencing

We isolated 22 novel *C. perfringens* strains from broilers and environmental samples from farms in Oxford, UK. Genomic DNA of these bacterial isolates was extracted using phenol-chloroform method as described previously [59]. Details of these isolates are given in Additional file 1: Table S1. Sequencing was performed at the Wellcome Trust Sanger Institute using Illumina HiSeq 2500 to generate 125 bp paired-end reads. Illumina reads are available in the European Nucleotide Archive under project PRJEB32760.

### Genome assembly and annotation

Broiler-related *C. perfringens* genome assemblies (RefSeq) and quality-trimmed sequencing reads (SRA) were retrieved from NCBI databases in May 2019 including available metadata (n=68). A total of 22 newly sequenced genomes were assembled in-house. All adapter-trimmed sequencing reads were used as input for MEGAHIT v1.1.1 [60]. Genome assembly was carried out using MEGAHIT options --k-min 27 --k-max 247 (for paired-end reads 2 x 250 bp), --k-min 27 --k-max 97 (for paired-end reads 2 x 125 bp), --no-mercy (specifically for generic assembly) and --min-contig-len 300. Over-fragmented draft genome assemblies were excluded from further computational analysis if >500 contigs (n=2). Assembly statistics were calculated using custom Perl script and all sequences were checked to have ANI >95% with respect to type strain ATCC13124 genome (Additional file 1: Table S2). All genomes were annotated using Prokka v1.13 with specific *Clostridium* genus (35 *Clostridium* species from NCBI RefSeq annotations) database with parameters --usegenus --mincontiglen 300 (Additional file 1: Table S3).

### Phylogenetic analysis, SNP detection, *in silico* virulence gene and plasmid detection

Annotated gff files were used as input for Roary v3.12.0 to construct pangenome with option -e -n to generate a core gene alignment via MAFFT, -s do not split paralogs, -i to define a gene at BLASTp 90% identity and -y to obtain gene inference [61]. A total of 20194 single nucleotide variants (315 715 site alignment from 1810 core genes) were called using snp-sites v2.3.3 [62]. We used the 20194 site-alignment to infer a ML phylogeny using RAxML v8.2.10 with GTR+ nucleotide substitution model at 100 permutations conducted for bootstrap convergence test [63]. The ML tree constructed was with the highest likelihood out of 5 independent runs (option -N 5). Pairwise SNP distances were calculated using snp-dists v0.2 [64]. ANI was computed using module pyani v0.2.7 [65]. R package *rhierBAPS* was used for phylogenetic clustering analysis to identify population structure [66].

Nucleotide sequence search was performed using ABRicate v0.8.11 on genome assemblies with coverage ≥90% and sequence identity ≥90% [67]. Toxin database was constructed as previously described [20] and collagen adhesin genes was detailed in Additional file 1: Table S4. Plasmids were predicted computationally using PlasmidSeeker v1.0 where sequence reads are available (k-mer coverage >80%), and ABRicate on all genome assemblies, with best-hit approach at query coverage threshold ≥70% and nucleotide identity ≥90% via custom database as detailed in Additional file 1: Table S5.

### Genome-wide gene association analysis

Scoary v1.6.11 was run to draw gene associations at default parameters [68]. Specificity cutoff was set at 80%, sensitivity at 100% to obtain 63 genes specifically associated with sub-lineage Vf isolates (Additional file 1: Table S6).

### 16S rRNA amplicon sequencing and analysis

Genomic DNA extraction of caecal samples was performed with FastDNA Spin Kit for Soil following manufacturer’s instructions and extending the bead-beating step to 3 min as described previously [26]. Extracted DNA was quantified and normalized to 5 ng/μl for all samples before subject to 16S rRNA Illumina MiSeq sequencing library preparation, amplifying V1 + V2 regions of the 16S rRNA gene as detailed in Additional file 1: Table S7 for the primer sequences. PCR amplification conditions were: 1 cycle of 94 °C for 3 min, followed by 25 cycles of 94 °C for 45 s, 55 °C for 15 s and 72 °C for 30 s. Libraries were sequenced on the Illumina MiSeq platform using a read length up to 2 × 300 bp.

Sequencing reads were analysed using OTU clustering methods via QIIME v1.9.1 using SILVA_132 as reference database to assign OTU by clustering at 97% similarity [69, 70]. Briefly, paired-end sequences were merged using PEAR, followed by quality filtering using split_libraries_fastq.py, chimera identification using identify_chimeric_seqs.py and chimera removal using filter_fasta.py [71]. OTU picking step was run using open reference approach pick_open_reference_otus.py which does not discard unassigned reads in the final output. BIOM output file was visualised on MEGAN6 [72]. Paired-end approach of taxa assignment using BLASTn as described previously [26].

Caecal contents were processed to generate an average of 116967 (range: 80 426-152 763) raw sequence reads per sample, with an average of 606 (range: 364-1559) OTUs assigned in each sample, clustering at 97% similarity (Additional file 1: Table S8). Rarefaction analysis supports the availability of sufficient sequence reads to achieve asymptotic based on rarefied reads (normalized to the lowest reads of all samples), i.e. optimal diversity of richness (range: 15-23 genera, 15-25 families) in each individual sample to represent each member of the microbiota (Additional file 2: Fig. S1).

LDA was performed using LEfSe via Galaxy server to identify significantly enriched taxa in the dataset. Alpha value for non-parametric Kruskal-Wallis test was set at 0.05 and threshold on LDA score at 2.0 for statistical significance. Graph was illustrated using the LEfSe plotting module [73].

R package *vegan* function *rarecurve* was used to draw rarefaction curves using rarefied reads (normalised to the lowest-read sample as implemented in MEGAN6). Diversity indices including Inverse Simpson index, Shannon index and Fisher index were computed using R package *vegan* function *diversity* [74].

### Statistical analysis and graphing

Venni 2.1 was used to analyse plasmid data [75]. Graphpad PRISM v6.0 was used for various statistical analyses, R package *ggplot2* was used for various plotting.

## Abbreviations

ANI: Average Nucleotide Identity;
LDA: Linear Discriminant Analysis;
ML: Maximum-likelihood;
NE: Necrotic Enteritis;
PCA: Principal Component Analysis;
PFO: Perfringolysin O;
SNE: Sub-clinical Necrotic Enteritis;
WGS: Whole Genome Sequencing

## Declarations

### Ethics approval

Not applicable

### Consent for publication

Not applicable

### Availability of data and materials

The key datasets analysed during the current study are available as follows:

1. Sequence reads for 22 newly-sequenced *Clostridium perfringens* strains were deposited under project accession number PRJEB32760.
2. 16S rRNA sequence reads of 11 caecal samples were deposited under project accession number PRJEB33036.
3. Phylogenetic tree aligned with metadata and virulence profiles is available in iTOL: https://itol.embl.de/tree/149155196252227531559222489

## Acknowledgments

This research was supported in part by the NBI Computing infrastructure for Science (CiS) group through the provision of a High-Performance Computing (HPC) Cluster. We also thank the sequencing team at Wellcome Trust Sanger Institute for genome sequencing.

## Authors’ contributions

RK and LJH designed the study. RK processed the sequencing data, performed the genomic and 16S rRNA amplicon analysis, and graphed the figures. SC provided essential assistance in genome assembly and genomic analysis. RK and LJH analysed the data and co-wrote the manuscript along with JB, RAD and GD. CL processed the caecum content samples and sequencing library. RK performed the full-length 16S rRNA PCR. JB collected the broiler faecal and caecal samples, isolated bacterial strains. RK and HB extracted genomic DNA from pure cultures for genome sequencing, which was supported by DP. All authors read and approved the final manuscript.

## Additional files

Additional file 1:

- File name: Additional_file_1.xlsx
- Title: Supplementary tables

Additional file 2:

- File name: Additional_file_2.docx
- Title: Supplementary figures

