## Supplementary figures and images for "Genomic analysis on broiler-associated *Clostridium perfringens* strains and caecal microbiome profiling reveals key factors linked to poultry Necrotic Enteritis"

### Additional file 2

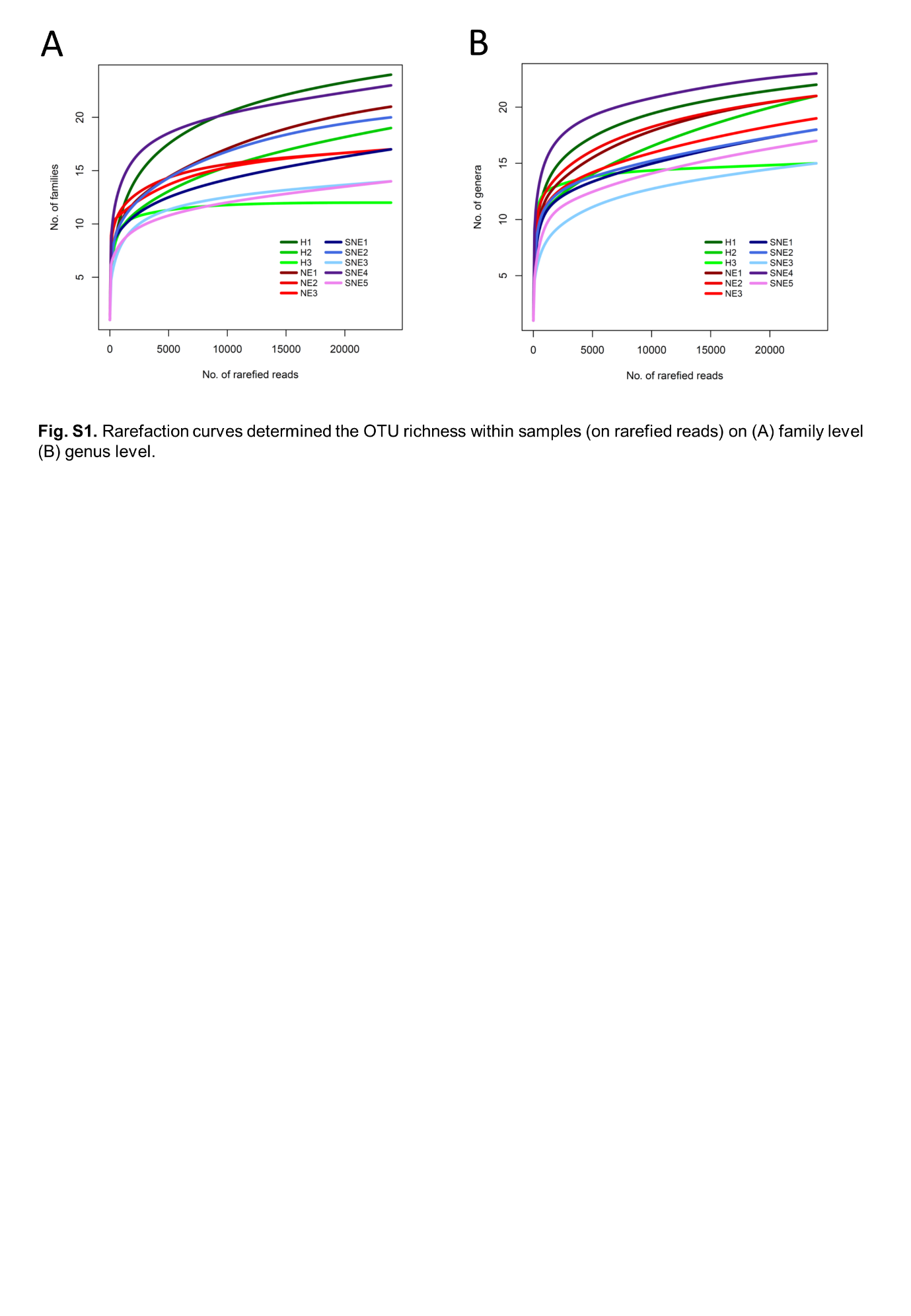

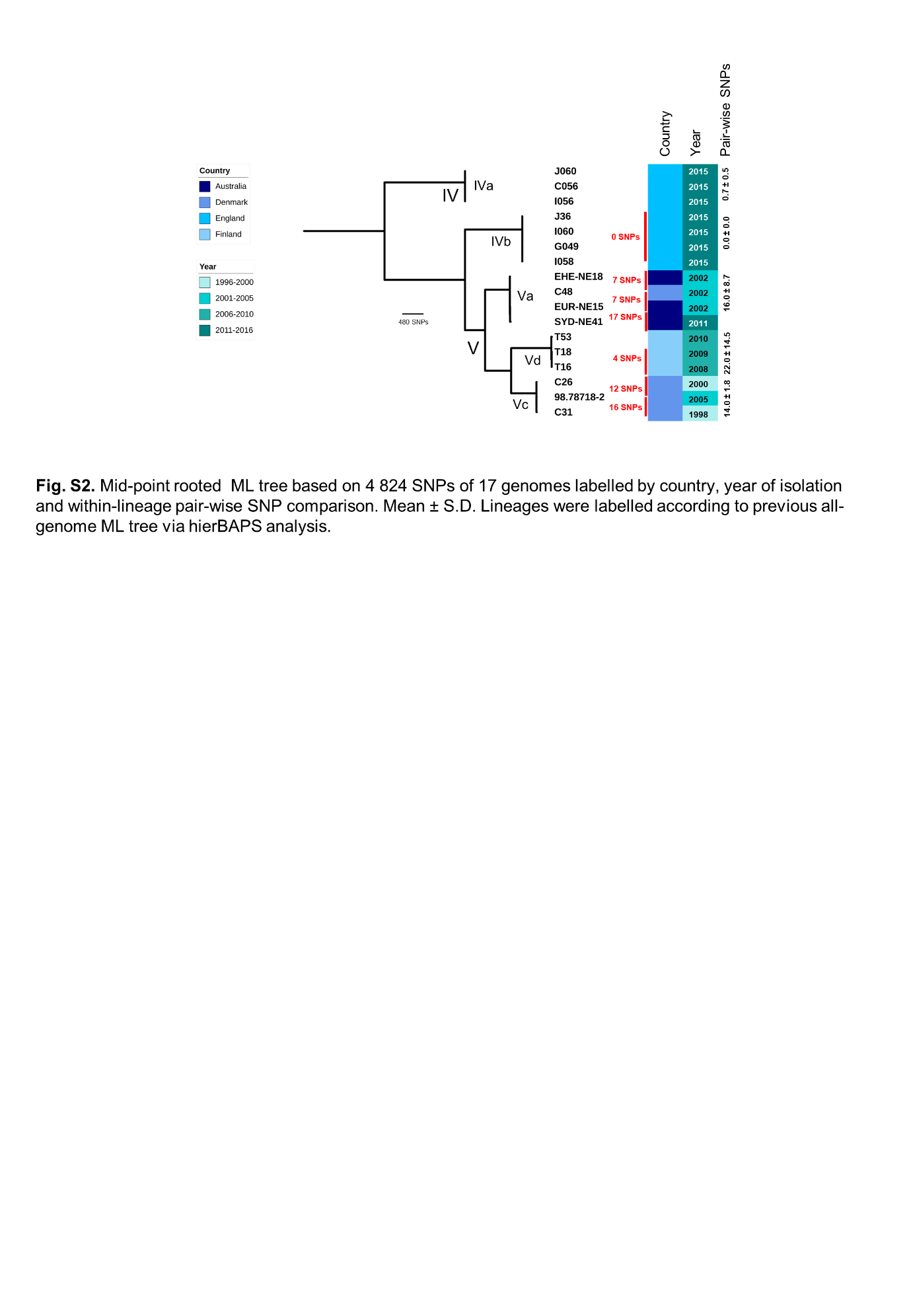

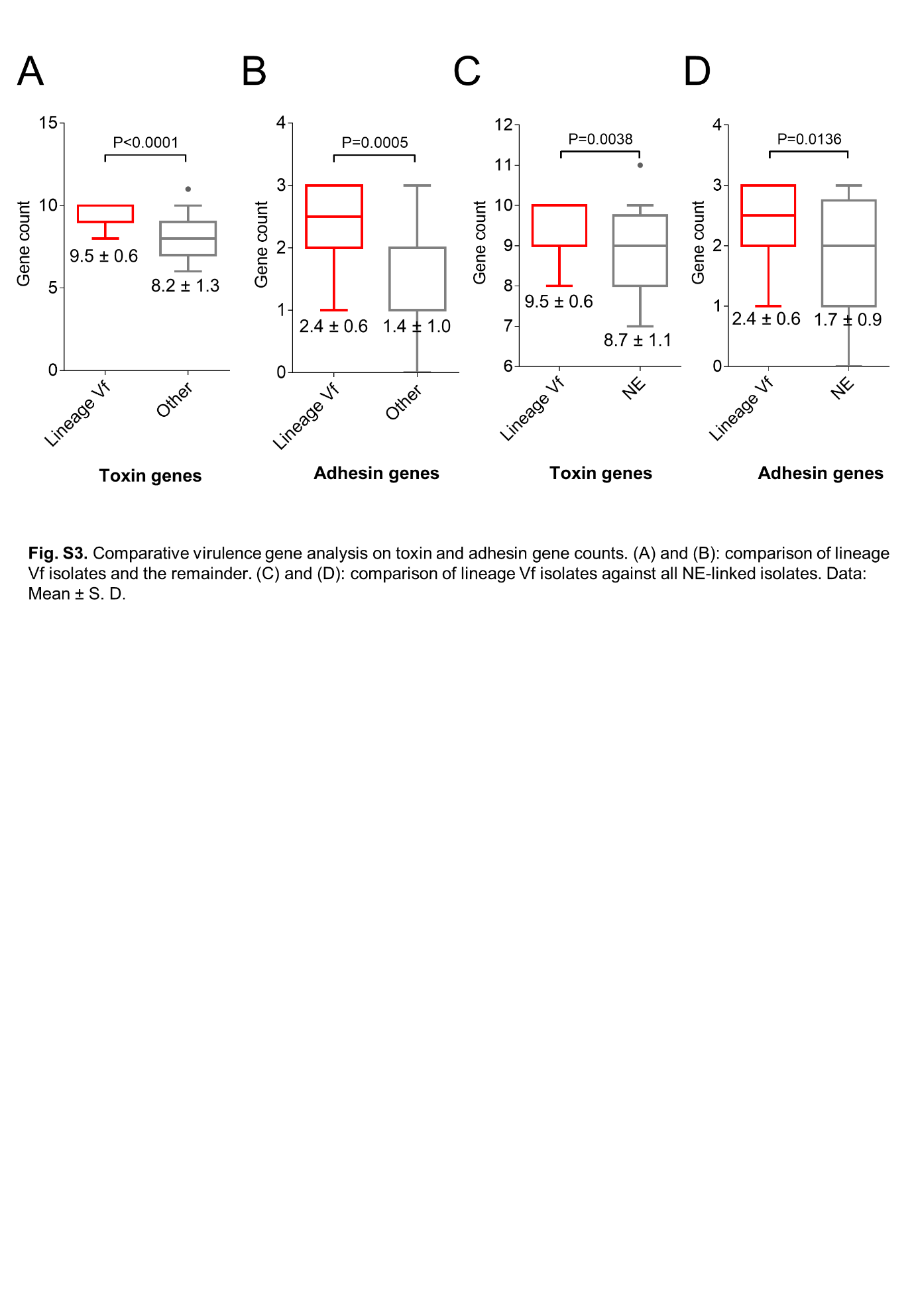

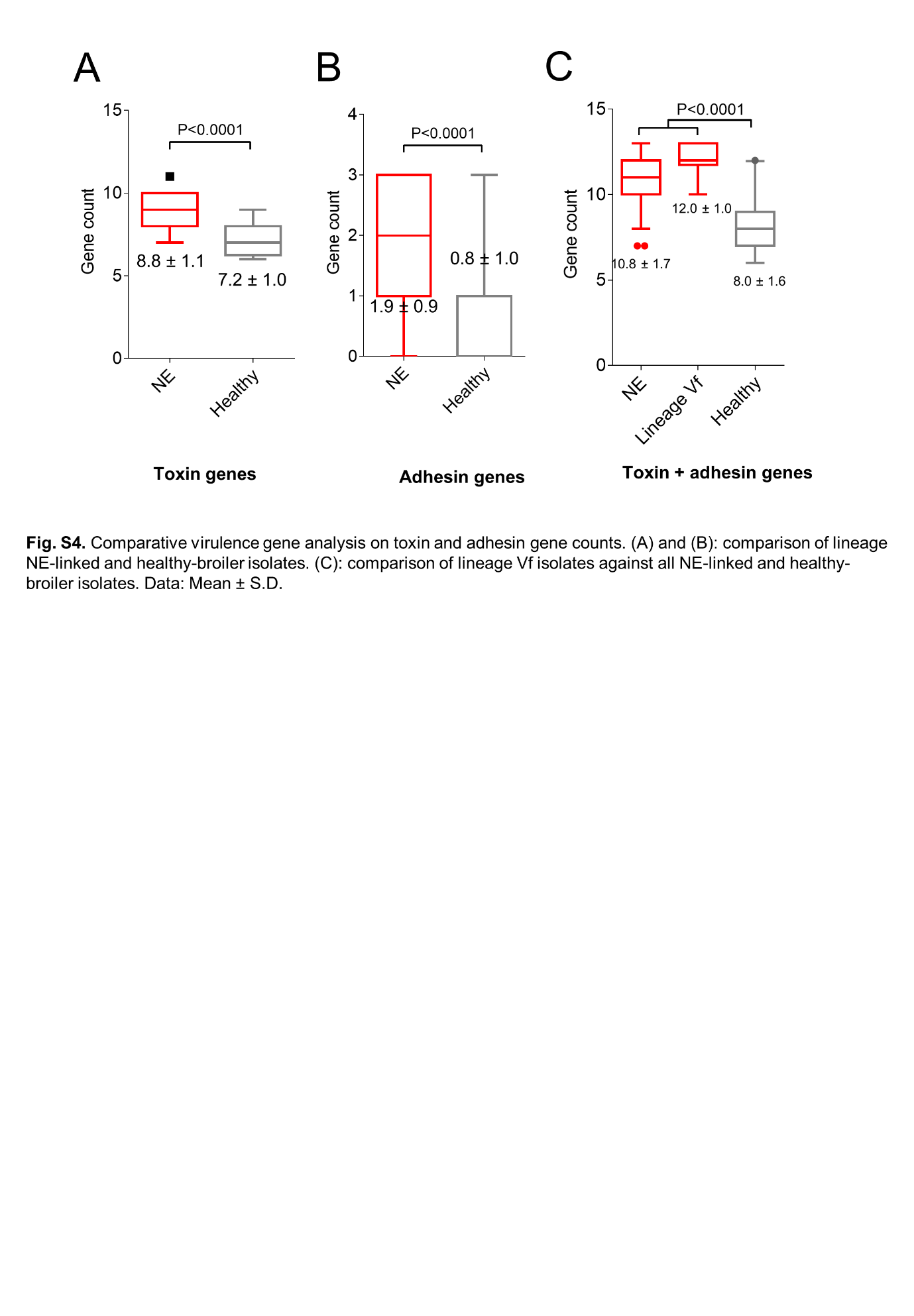

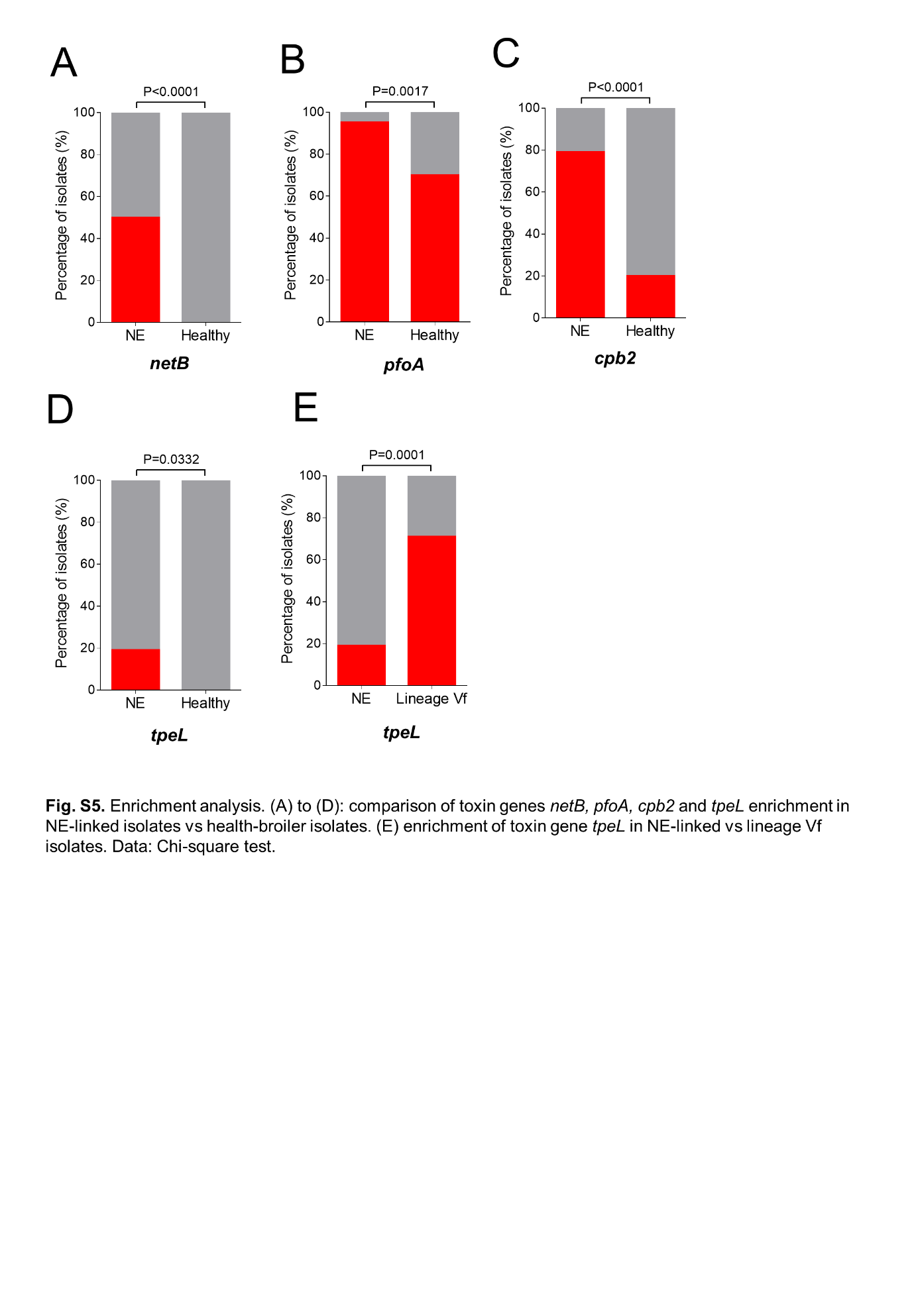

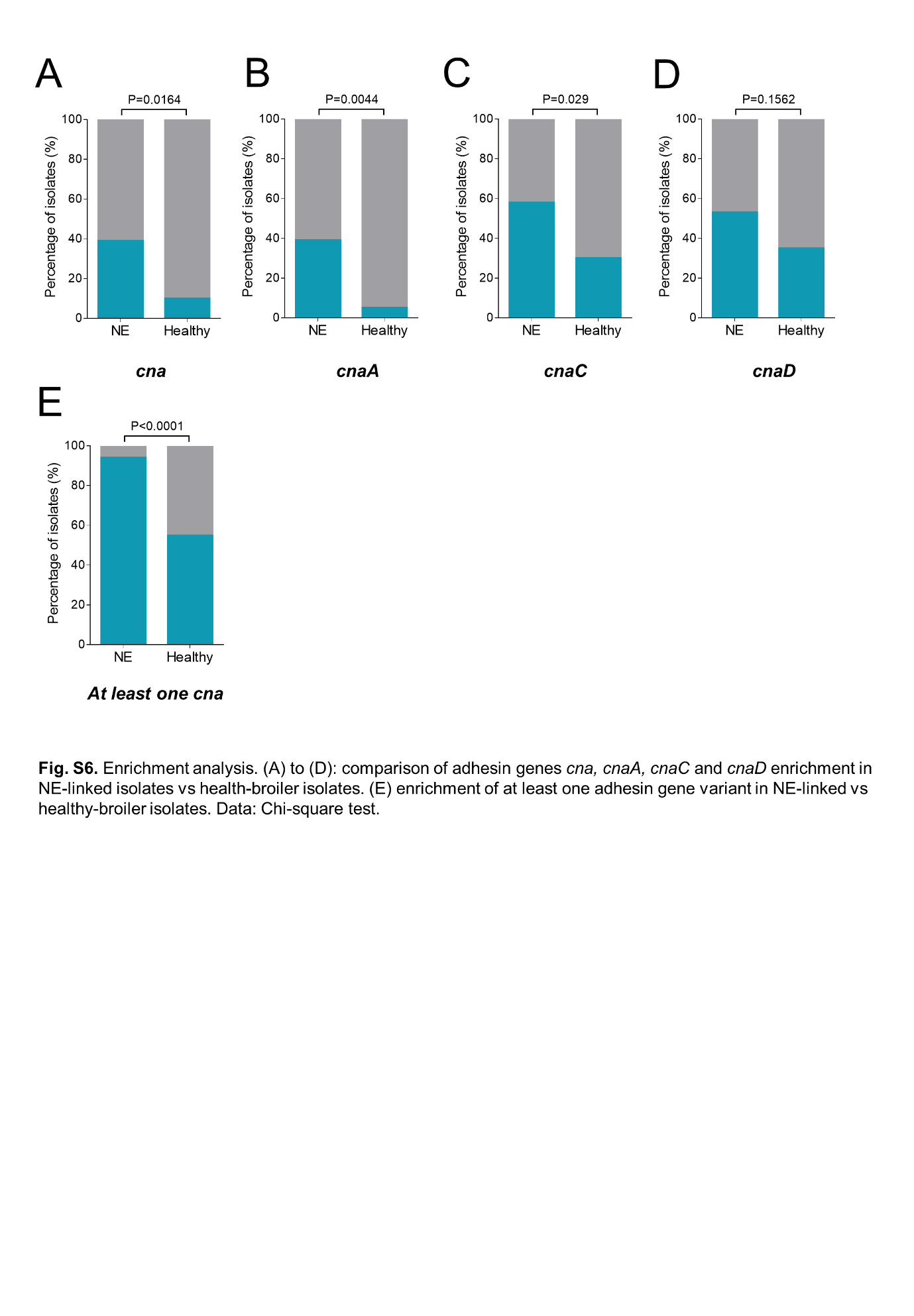

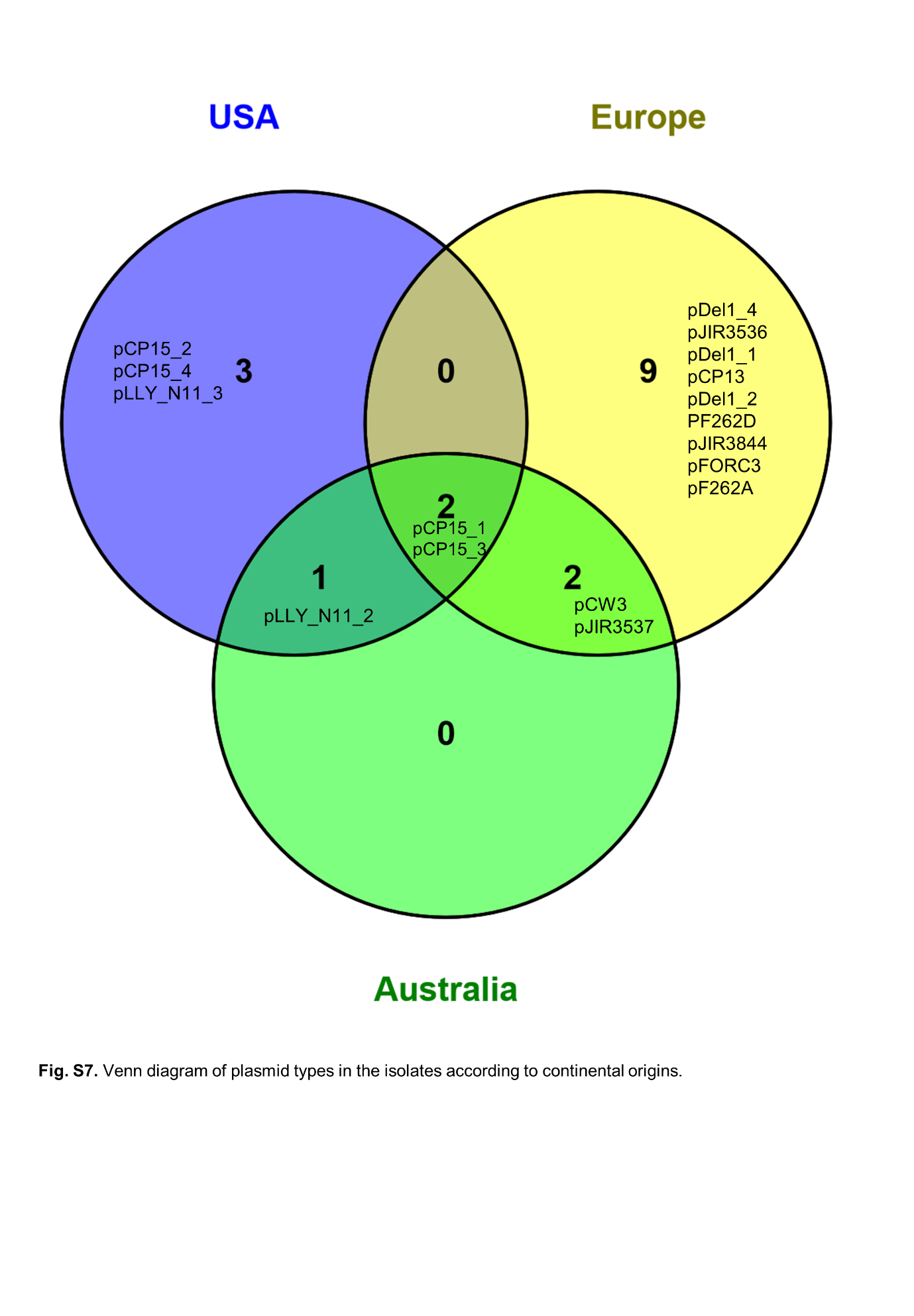

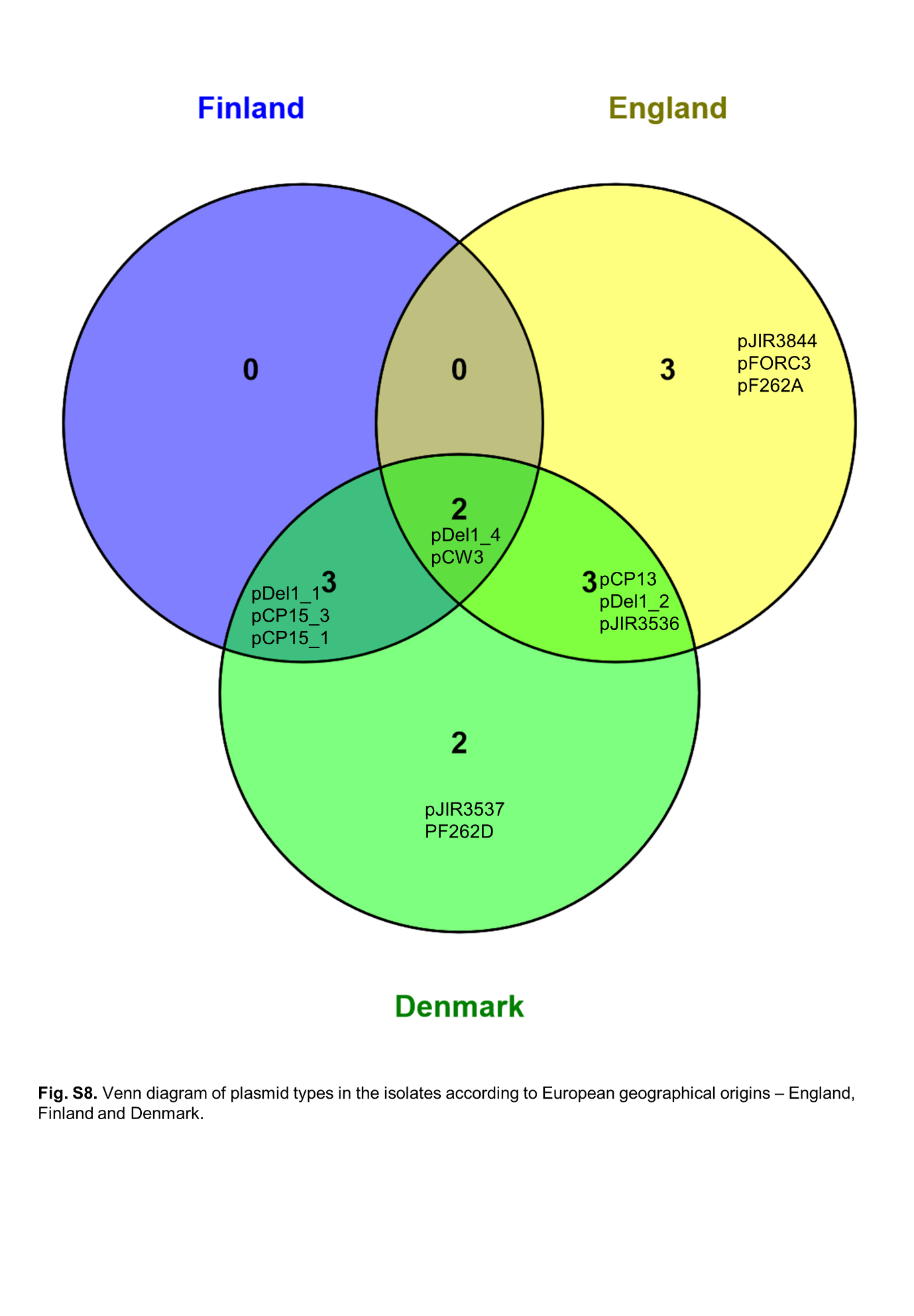

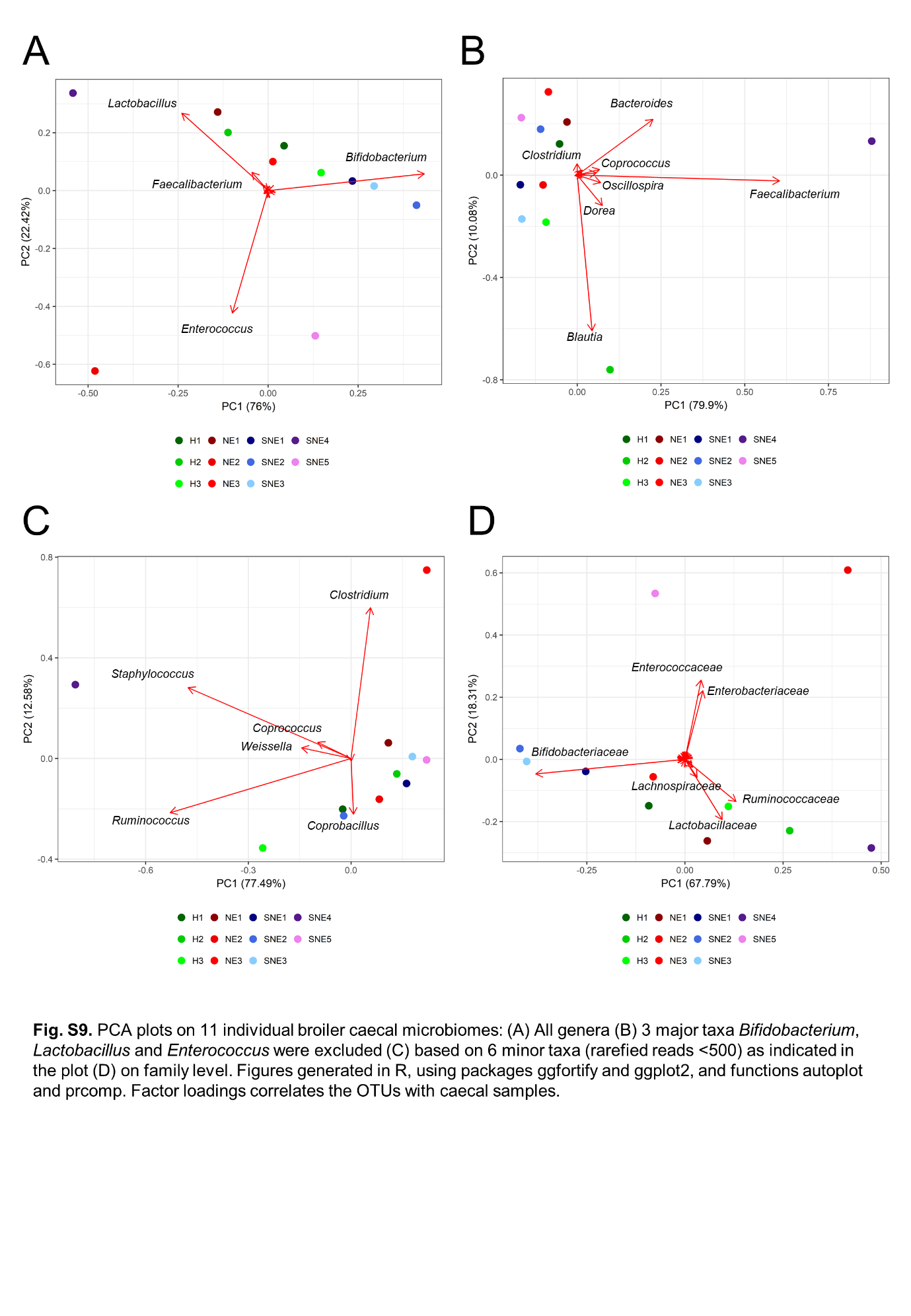

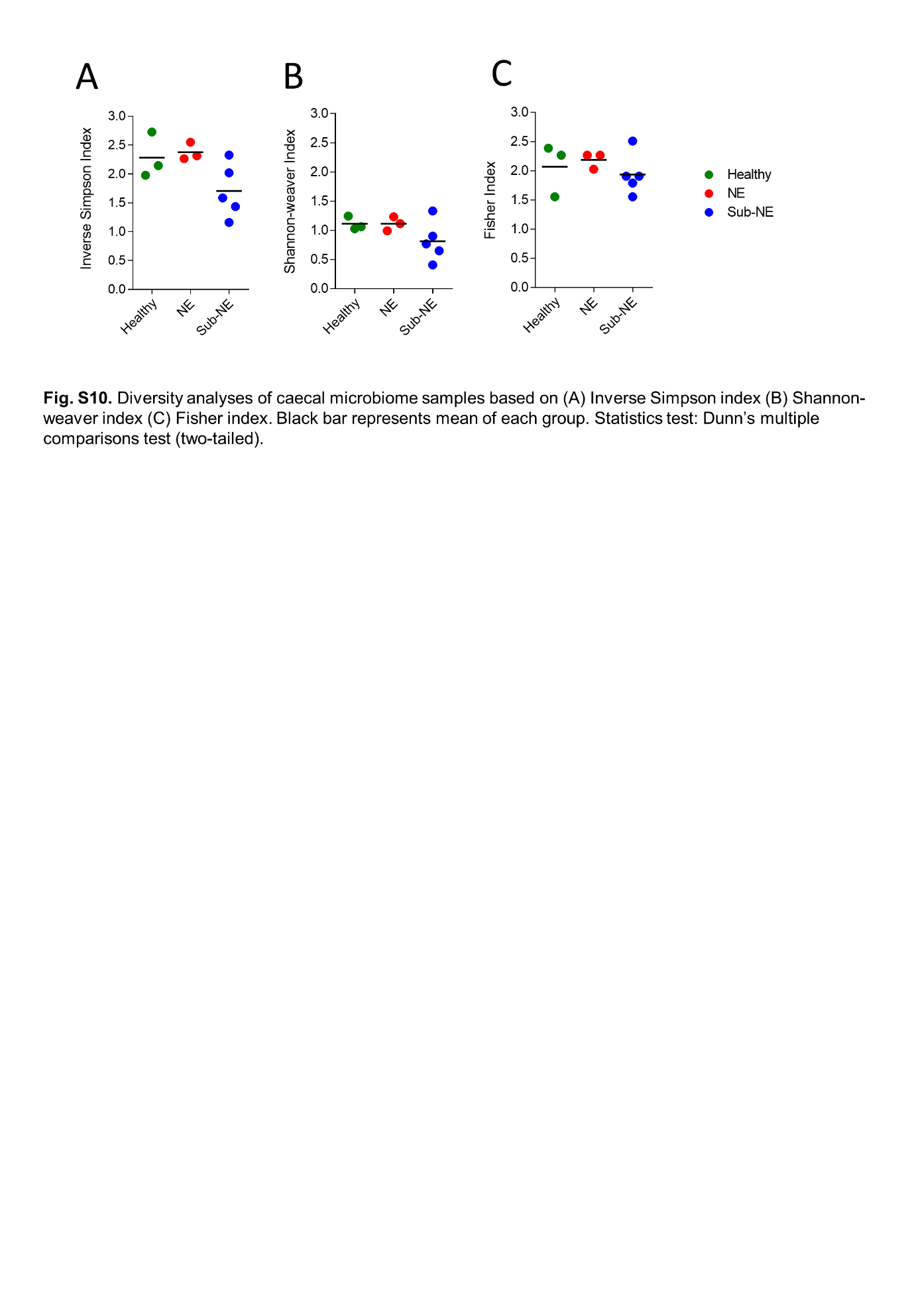

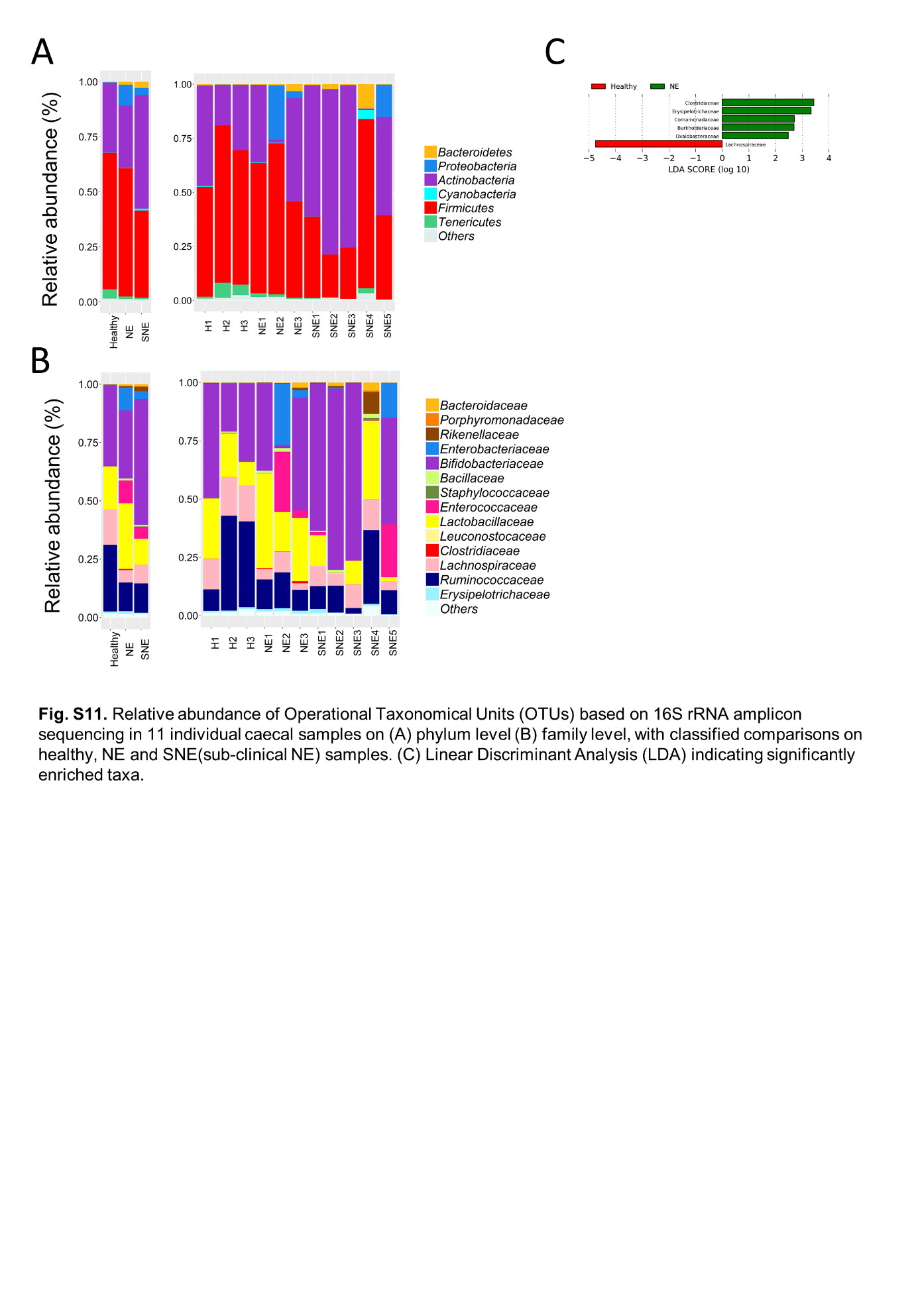

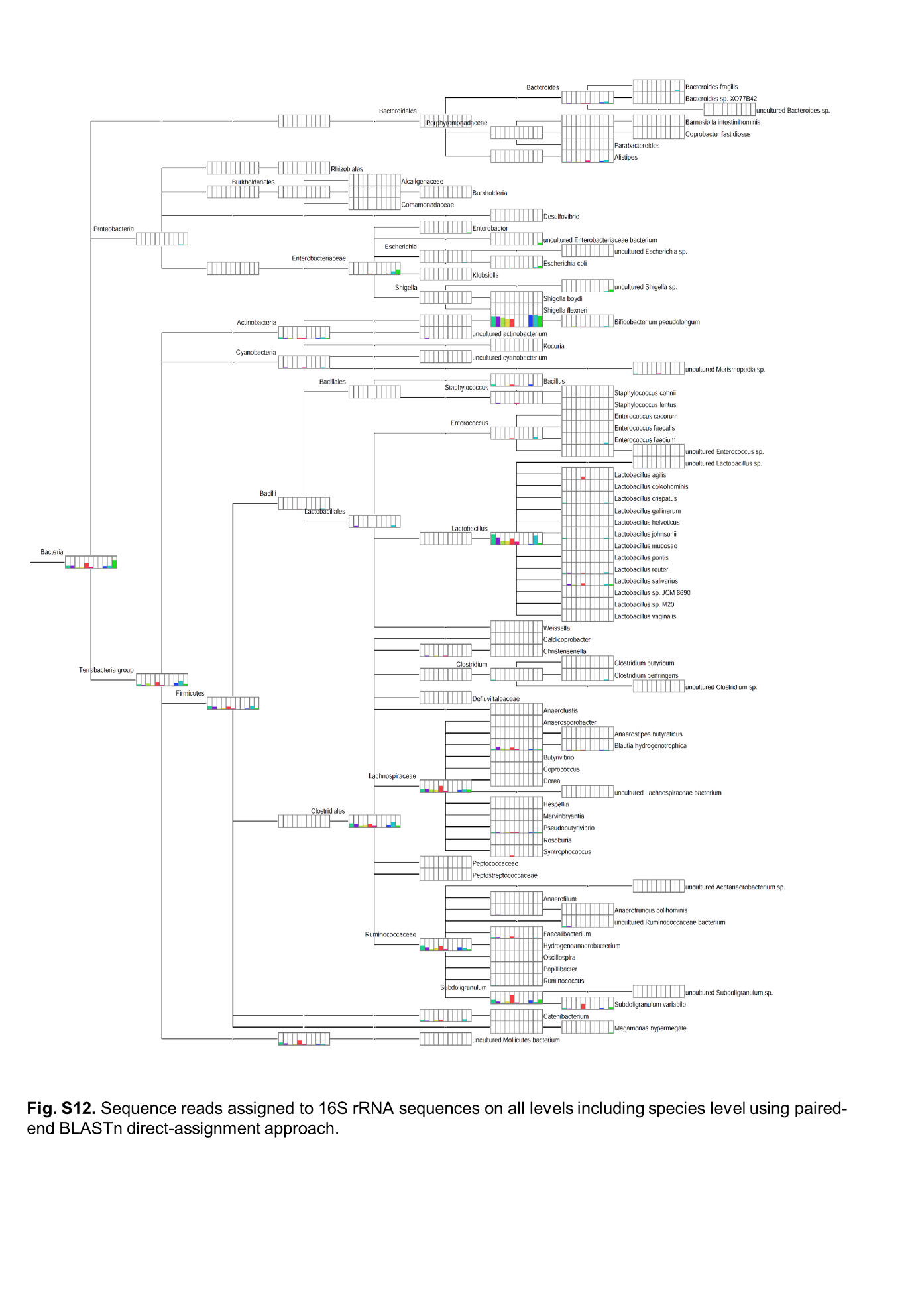
